# Global and selective effects of auditory attention on arousal: insights from pupil dilation

**DOI:** 10.1101/2024.11.19.624303

**Authors:** Aurélie Grandjean, Roxane S. Hoyer, Anne Mathieu, Anne Caclin, Annie Moulin, Aurélie Bidet-Caulet

## Abstract

Theoretical models of attention propose that norepinephrine (NE) can induce both a global boost of arousal and selective amplification of high-priority stimuli, yet few tasks have tested these dual effects in humans. Here, we used pupillometry in an auditory detection task, the Competitive Attention Test (CAT), previously performed in large cohort studies, to examine how task engagement (active vs. passive) and stimulus relevance (informative vs. uninformative cues) modulate arousal. Results showed that both relevant and irrelevant sounds elicited larger pupil dilation under active conditions, indicating a global arousal effect. Crucially, only relevant sounds benefited from an additional dilation when preceded by an informative cue, demonstrating a selective arousal mechanism associated to top-down attention. These findings illustrate the NE’s dual role in boosting overall alertness while selectively enhancing high-priority stimuli. Beyond theoretical implications, this work highlights that the CAT captures measurable arousal components, reinforcing its utility for clinical assessments of attention-arousal clinical disruptions.

## Introduction

### Arousal and Attention: theoretical models

Efficient attention abilities and optimal level of arousal are critical to adapt to the ever-changing environment (Kahneman, 1973; Howels et al., 2012). Arousal refers to a physiological state of alertness or reactivity (Broadbent, 1971; Kahneman, 1973; Coull, 1998; Näätänen, 1992; Sturm & Willmes, 2001), and more recently, to a general state of cortical excitation driven by the activity of the locus coeruleus - norepinephrine (LC-NE) system (Aston-Jones & Cohen, 2005; Nieuwenhuis et al., 2011). The LC is a brainstem neuromodulatory nucleus responsible for most of the NE release in the brain through widespread projections to the entire neocortex (Berridge & Waterhouse, 2003). A moderate level of arousal has been shown to result in improved cognitive performance, whilst reduced and enhanced levels of arousal lead to distractibility (Yerkes & Dodson, 1908; Easterbrook, 1959). Attention enables us to prioritize and focus on relevant information, while still processing irrelevant stimuli, allowing us to stay aware of – but yet not distracted by – unexpected events in our environment. Achieving this requires a good balance between a “global” readiness to respond (i.e., arousal) and a “selective” focusing mechanism (i.e., voluntary attention) to filter out distracting stimuli. Attention is assumed to control resources allocation to task-relevant events, while task engagement refers to the sustained allocation of cognitive resources to a task, thus elevating arousal (Tops and Boksem, 2010). Importantly, dysregulation in either process is linked to conditions such as attention deficit disorder or dementia, underlining the current clinical need to measure how arousal and attention interact (for examples, Malhotra, 2019; Ohm et al., 2020). Understanding the dynamic interplay between arousal and attention is indeed essential for precise diagnosis when attention is altered as part of clinical condition, and for developing tailored therapeutic interventions.

Multiple attention theories suggest a dynamic interplay between arousal and selective attentional processes. For instance, Kahneman’s (1973) capacity model posits that phasic (i.e., transient) and tonic (i.e., sustained) arousal jointly shape the pool of available attentional resources, while the orienting response to novel stimuli reflects an involuntary rise in arousal (see also Näätänen (1992) for a similar proposal). Later, Broadbent (1971) and Eysenck (1982) differentiated between a “passive”, low-level system, and an “active”, higher-level cognitive system, each regulating arousal differently. They proposed that the passive system regulating arousal during task performance would itself be regulated by the active system to optimize task performance. Importantly, recent theories suggested that arousal does more than maintain overall alertness: it would also refine *selective* attention mechanisms (Dahl et al., 2022). Indeed, through NE release in the brain, arousal can amplify strong task-relevant signals while suppressing weak or random activity, thereby selectively facilitating the processing of relevant stimuli (Mather et al. 2016). However, to date, we lack a unified account of how top-down attention might engage both a global boost in arousal (task engagement) and a more targeted enhancement for high-priority events (e.g., validly cued targets).

### Arousal and attention: brain and behavioral findings

Anatomical brain connections further support the interplay between arousal and attention. Experimental studies, mostly in rodents and monkeys, showed widespread projections from the LC-NE system to the cerebral cortex, the cerebellum, and subcortical structures (thalamus, amygdala) (Berridge & Waterhouse, 2003). Conversely, attention control is supported by brain networks, including frontal and parietal regions (Corbetta & Shulman, 2002; Posner & Peterson, 1990), with descending connections toward the locus coeruleus (Corbetta et al., 2008; Schwarz et al., 2015; Breton-Provencher & Sur, 2019; Totah et al., 2021). However, the functional interplay between arousal and attention remains poorly understood as only a few non-invasive methods allow to measure deep brain activity within structures such as the LC in humans.

Physiological markers such as heart rate, skin conductance, and pupil dilation are considered as indirect measures of the arousal level (Berntson, 2007; Wang et al., 2018). Among them, pupil dilation can be considered the most sensitive method to investigate the relationship between arousal and attention during task performance, as pupil dilation has previously been related to the LC-NE activity in animal models (Aston-Jones & Cohen, 2005; Gilzenrat et al., 2010; Joshi et al., 2016). Specifically, baseline changes in pupil size are thought to reflect tonic activity in the LC, associated with tonic arousal, which has extensively been associated with sustained alertness and task engagement (Aston-Jones & Cohen, 2005; Gilzenrat et al., 2010; Jepma & Nieuwenhuis, 2011; Murphy et al., 2014; Unsworth & Robison, 2016, 2018; Konishi et al., 2017). Furthermore, the stimulus-related pupil response (PR) is thought to reflect the LC phasic activity and phasic arousal (Murphy et al., 2011; Wetzel et al., 2016). An increasing number of studies has investigated the relationship between arousal and attention using pupil dilation in the auditory modality (for review Zekveld 2018). Several studies showed that the PR is larger to deviant or novel sounds compared to standard sounds in passive oddball paradigms (Friedman et al., 1973; Steiner & Barry, 2011; Hong et al., 2014; Korn & Bach, 2016; Wetzel et al., 2016; Marois et al., 2018; Marois & Vachon, 2018; Widmann et al., 2018; Bonmassar et al., 2020; Liao et al., 2016), in line with an increase in phasic arousal.

Therefore, measuring pupil dilation during auditory attention tasks presents a promising approach to studying the interplay between arousal and attention. In a first body of experiments, the effects of auditory attention on the PR have been investigated by manipulating task demand, from passive to the most complex tasks. Asking participants to actively perform a task (by focusing and providing a motor response) or to passively follow the presentation of the same stimuli can be considered as the experimental paradigm allowing the maximum contrast in terms of amount of attention required. The mean PR to novel stimuli seemed to be larger in active than passive oddball conditions (though not statistically tested, Liao et al., 2016). By manipulating the task difficulty, some studies showed that the pupil dilation was larger for the most complex tasks (increased difficulty of tone discrimination: Kahneman and Beatty, 1967; words-in-noise identification task: Kramer et al., 2012). By manipulating attentional load, Lisi et al. (2015) showed a larger pupil dilation during multitasking than in a single task condition. Moreover, in several experiments using Posner visual cueing tasks to manipulate voluntary attention orienting, the amplitude of PR toward targets increased (Dragone et al. 2018) or remained unchanged (Aminihajibashi et al. 2020) with increasing cue predictiveness. These studies, which vary the overall amount of attentional resources required for task performance, suggest that enhancing task engagement results in increased arousal.

Effects of auditory attention on the PR could also be investigated by specifically manipulating stimulus relevance, i.e., exploring how attention facilitates or inhibits the PR to task-relevant and -irrelevant stimuli. To date, only Liao et al. (2016) have investigated this question by manipulating attention during an audio-visual oddball paradigm: the PR amplitude observed in response to deviant sounds did not differ according to task-relevance (attention directed to the auditory or visual modality). These limited findings raise questions about how the interplay between arousal and attention is mediated.

### The present study

According to the Glutamate Amplified Noradrenergic Effects (GANE) model (Mather et al., 2016), as well as recent work by Dahl et al. (2022), NE not only globally heightens arousal but also amplifies the processing of high-priority, task-relevant information. This dual, “selective amplifier” role of NE has yet to be fully tested in a paradigm that manipulates both task engagement (active vs. passive) and stimulus relevance (informative vs. uninformative cues). Consequently, we employed pupillometry in a recently attention test, the Competitive Attention Test, to examine precisely how NE-driven arousal interacts with voluntary attention orienting.

The Competitive Attention Test (CAT), developed by Bidet-Caulet and collaborators (2015), is an auditory detection paradigm, enabling the investigation of arousal responses to both relevant and unexpected irrelevant events (Figure 1). In this paradigm, voluntary attention orienting is manipulated by varying the predictive value (informative vs. uninformative) of a visual cue preceding a target sound to be detected. Auditory distraction and phasic arousal are triggered by unpredictable task-irrelevant sounds (distracting sounds) presented at different times between the cue and the target. In the present study, we also manipulated participants’ level of task engagement by comparing an active detection condition with passive exposure to the same stimuli. The arousal level was indirectly assessed by measuring pupil dilation (Gilzenrat et al., 2010; Nieuwenhuis et al., 2011; Joshi et al., 2016).

**Figure 1.**
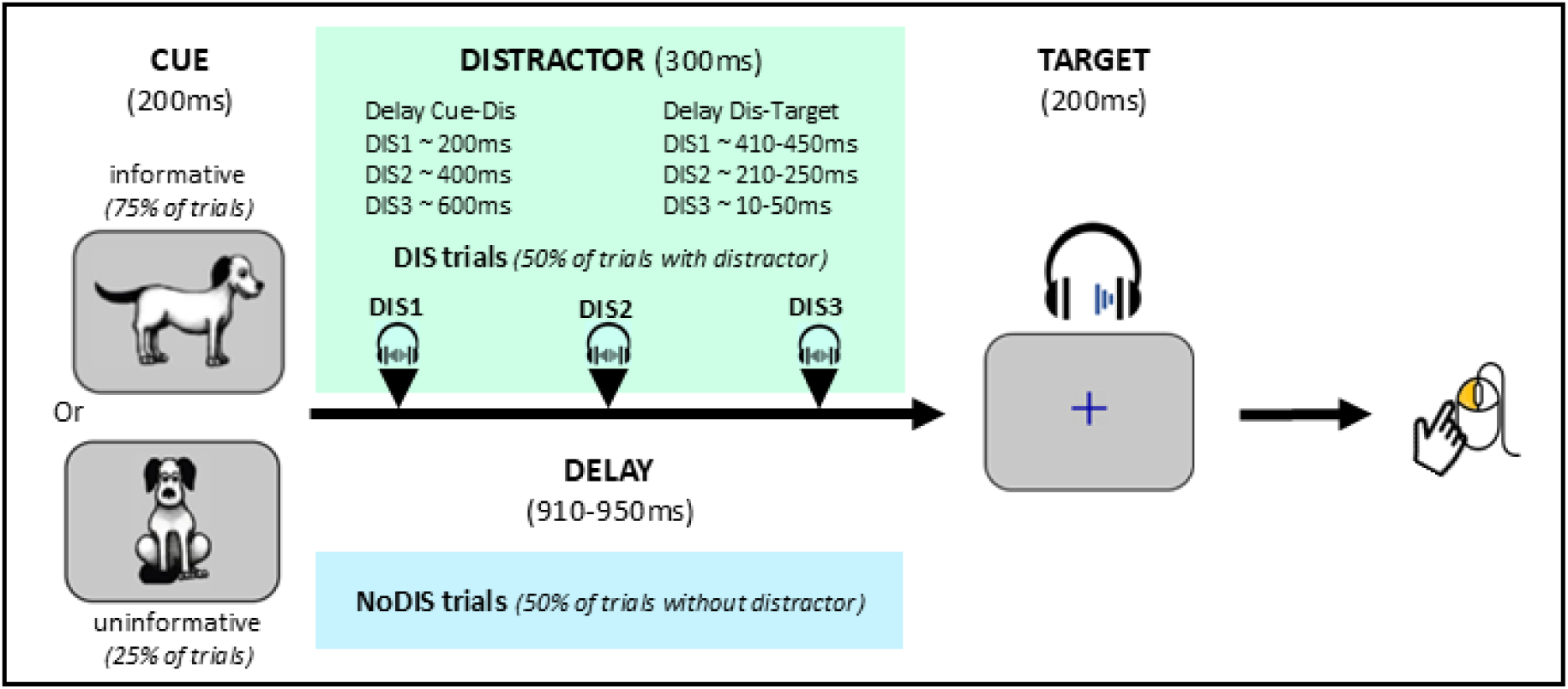
Protocol for active trials. All trials started with a visual cue (200ms duration) and contained a monaural target sound presented 910-950ms after cue offset. Subjects were asked to press the mouse button as fast as possible when they heard the target sound (a dog bark). In trials without distractor (NoDIS trials, 50%, blue box), only the cue and the target were presented. In trials with distractors (DIS trials, 50%, green box), a binaural distracting sound (300ms duration, symbolized by black triangles) was presented during the delay. The distracting sound could equiprobably appear in three different time periods before the target onset: 410-450ms (DIS1), 210-250ms (DIS2), and 10-50 ms (DIS3) (see corresponding cue-distractor delays in the figure). In 75% of the (noDIS and DIS) trials, a dog facing left or right indicated in which ear (left or right) the target sound would be played (informative cue). In the other 25% of the trials, a dog facing front did not provide any indication in which ear the target sound would be played (uninformative cue).

According to the current framework (e.g., GANE model), we hypothesized that *task engagement* (active vs. passive) would elicit larger PR to *all* events, reflecting a global arousal increase; whereas *voluntary attention orienting* (informative vs. uninformative cues) would further *selectively* amplify the PR for task-relevant over irrelevant stimuli.

## Method

### Participants

Twenty-three paid healthy adults participated in the experiment. Three participants were excluded due to excessive missing data in the pupillometry recordings or after preprocessing, leading to insufficient number of trials for the subsequent analyses. Therefore, 20 participants (mean age ± standard deviation: 23.3 ± 3.8 years old; 12 female, 17 right-handed) were included in the analyses. This sample size is similar to previous studies using similar paradigms (Bidet-Caulet et al.,2017; ElShafei et al., 2018) and pupillometry (Preuschoff et al., 2011; Koelewijn et al., 2014; Lisi et al., 2015; Dragone et al., 2018).

All participants were free from any neurological or psychiatric disorder, had normal hearing and normal or corrected-to-normal vision, and did not take any substance (alcohol, drug, or medication) affecting the central nervous system during the 24 hours preceding the testing session. Finally, participants were instructed not to consume coffee or energy drinks for at least three hours before the testing session. All participants gave written informed consent. This study was conducted according to the Helsinki Declaration, Convention of the Council of Europe on Human Rights and Biomedicine, and the experimental paradigm was approved by a French ethics committee (Comité de Protection des Personnes Sud-Est IV, number 11/90, authorization B11291-10).

### Stimuli

Here we used the same stimuli and trial structure than in previous study employing the CAT (Hoyer et al., 2021 & 2023a). All trials started with a visual cue (200ms duration) depicting a dog facing left, right or front, centrally presented on a grey background screen. Each trial also included a monaural target sound (200ms duration, 5ms rise-time and 5ms fall-time, 15 dB SL, around 43 dBA) corresponding to a dog bark played 910-950ms after cue offset. In trials without distractor (NoDIS trials, 50%), only the cue and the target were presented. In trials with distractor (DIS trials, 50%), a binaural distracting sound (300ms duration, 35 dB SL, around 61 dBA) was played during the cue-target delay. The distracting sound could appear with equal probability in three different time periods before the target onset: 410-450ms (DIS1), 210-250ms (DIS2), and 10-50 ms (DIS3) (Figure1).

In 75% of the (noDIS and DIS) trials, a dog facing left (37.5%) or right (37.5%) indicated in which ear (left or right) the target sound would be played (informative cue). In the other 25% of the trials, a dog facing front did not provide any indication in which ear (left: 12.5% or right: 12.5%) the target sound would be played (uninformative cue).

A total of eighteen RWICEdifferent sounds were used as distracting sounds (8 different clock-alarms, 8 different phone ringtones, 2 different door-bells). These sounds loudness was subjectively equalized by two listeners (Masson & Bidet-Caulet, 2019).

The inter-trial-interval duration (post target offset) was jittered between 3700 and 3900ms.

### Task

The stimuli were presented to the participants either passively (passive condition, see below) or actively (Active Audio-Visual condition). Although the different stimuli serve as cues, targets, or distracting sounds only in the active condition, the same terminology is used for clarity in the passive conditions.

In the Active Audio-Visual condition (AAV), participants were instructed to perform a detection task by pressing a mouse button as fast as possible when they heard the target sound (i.e., the dog bark) and to ignore the distracting sounds. The AAV condition included 3 blocks of 48 trials (total with informative cue: 54 NoDIS, 18 DIS1, 18 DIS2, and 18 DIS3 trials; total with uninformative cue: 18 NoDIS, 6 DIS1, 6 DIS2 and 6 DIS3 trials). Each distracting sound was played four times across the 3 blocks, but no more than twice during each block to minimize habituation.

In the Passive Audio-Visual condition (PAV), participants were instructed to passively watch and hear the same visual and auditory stimuli as in AAV, without providing any response. The PAV condition included 1 block of 48 trials (total with informative cue: 18 NoDIS, 6 DIS1, 6 DIS2 and 6 DIS3 trials; total with uninformative cue: 6 NoDIS, 2 DIS1, 2 DIS2, and 2 DIS3 trials).

Two additional passive blocks were added as control conditions. In the Passive Visual (PV) condition, only the visual cue was presented, in 24 trials (18 with informative cues and 6 with uninformative cues). Participants were instructed to passively watch the visual stimuli. In the Passive Auditory (PA) condition, only distracting and target sounds were presented, in 48 trials (24 NoDIS, 8 DIS1, 8 DIS2, and 8 DIS3 trials). Participants were instructed to passively listen to the sounds. Data from these control conditions are presented in the supplementary data section (SupFigure1).

### Procedure

#### Behavior

Participants were seated in a comfortable armchair at a 70-cm distance from a computer screen. All stimuli were delivered using Presentation software (Neurobehavioral Systems, Albany, CA, USA), which was also used to record the timing of behavioral responses. Sounds were delivered through circumaural headphones (Sennheiser 280HD). First, participants performed the PV block. Second, the auditory threshold was determined for the target sound, in each ear, for each participant using the Bekesy tracking method, later allowing to present the target sound at 15 dB SL and the distractor at 30 dB SL. Third, participants performed the PA and PAV blocks. Finally, they performed a short AAV training (10 trials), and three AAV blocks.

All participants performed the experimental blocks in the same order (PV, PA, PAV, AAV), so that the active condition did not contaminate the passive ones. Trials within blocks were pseudo-randomized differently for each participant to limit sequence effects and to compare responses to acoustically matched sounds across participants. Thus, across participants, the same distracting sounds were played for each distractor occurrence condition (DIS1, DIS2, or DIS3).

The total duration of the experiment was 1h.

#### Pupillometry

The pupil size of one eye (right: 13; left: 7) was recorded during each block using an eye-tracking system (Eyelink 1000) in arbitrary units (au) in a room with controlled luminosity. The illumination at proximity of the participants was around 7.5 lux. Participants were placed at a 79 cm distance from the eye-tracker which was set up in a remote mode at a sampling rate of 500 Hz. Each block started with a five-point eye-tracker calibration followed by a five-point validation procedure. Participants were instructed to blink naturally but minimize eye movements during the whole recordings. They were instructed to keep their gaze fixed on a blue cross, which was displayed continuously, except during the cue presentation.

### Data preprocessing

Data preprocessing was performed using the software package for electrophysiological analysis (ELAN Pack) developed at the Lyon Neuroscience Research Center (; Aguera et al. 2011) and custom MATLAB and Python programs.

#### Behavioral data

Reaction times (RT) limits after target onset for correct detections were chosen based on previous studies (Hoyer at al., 2021, 2023a, 2023b). The RT lower limit was 150ms and the RT higher limit was the mean of RT plus 2 standard deviations in each subject. A button-press before the RT lower limit was considered as a premature response. A trial with a button-press after the upper limit was considered as a late response. A trial with no button-press before the next cue onset was considered as a miss. The median of all RTs to targets was calculated for each subject.

#### Pupil data

Full blinks were detected and marked by the eye tracker. Segments with blinks or missing data were interpolated with linear interpolation. Trials with more than 60% of interpolated data were removed from further analysis (Seropian et al., 2022). Trials with premature response or missed trials were also excluded. The final mean percentage of rejected trials was (mean ± standard deviation) 2.3 ± 4.4 % in AAV, 1.8 ± 4.3 % in PAV, 0.6 ± 1.2% in PA & 2.1 ± 6.1 in PV. Participants with less than 20 remaining trials per condition were excluded from the subsequent analyses. Finally, data were low-pass filtered using a bi-directional Butterworth filter with an 80Hz cut-off.

Pupil responses (PR) were analyzed locked to either the cue or the distractor events.

Cue-locked PR was baseline-corrected by subtracting the 250ms period before cue onset.

Concerning DIS-locked PR, for task and cue effects, first a baseline subtraction was used, with a 250ms period pre-dis onset. Second, for each distractor onset time-range, “surrogate DIS-locked PR” were created in the NoDIS trials (also baseline corrected) and subtracted from the actual DIS-locked responses. The obtained DIS-locked-corrected PR to the distractor was thus clear of cue-related activity. For distractor position effects, only a baseline subtraction was used, with a 250ms period pre-dis onset for post-dis pupil size analysis and with a 250ms period pre-cue onset for pre-dis pupil size analysis.

### Statistical analysis

Frequentist statistical analyses were conducted using packages rstatix, emmeans, and PMCMRplus of R (version 4.2.2). Shapiro-Wilk test was used to test data normality. When data were found normally distributed, parametric analyses were performed; otherwise, non-parametric analyses were chosen.

#### Behavioral data (AAV condition)

Median RTs were submitted to a repeated-measure ANOVA with cue (two levels: informative, uninformative) and distractor (four levels: NoDIS, DIS1, DIS2, DIS3) as within-participant factors (Table 1.A, B). *F* values, probability levels and generalized eta squared (η²G) for effect sizes are provided. ηG2 values, representing the proportion of total variance explained by a factor, generally range from 0 to 1. According to Cohen’s interpretation guidelines (Cohen, 1988), the closer the value of ηG2 is to 1, the greater the effect of the factor. Conversely, the closer it is to 0, the weaker the effect. Significant main effects obtained with ANOVAs were further examined with pairwise paired t-test comparisons and the Bonferroni multiple testing correction method.

**Table 1.**
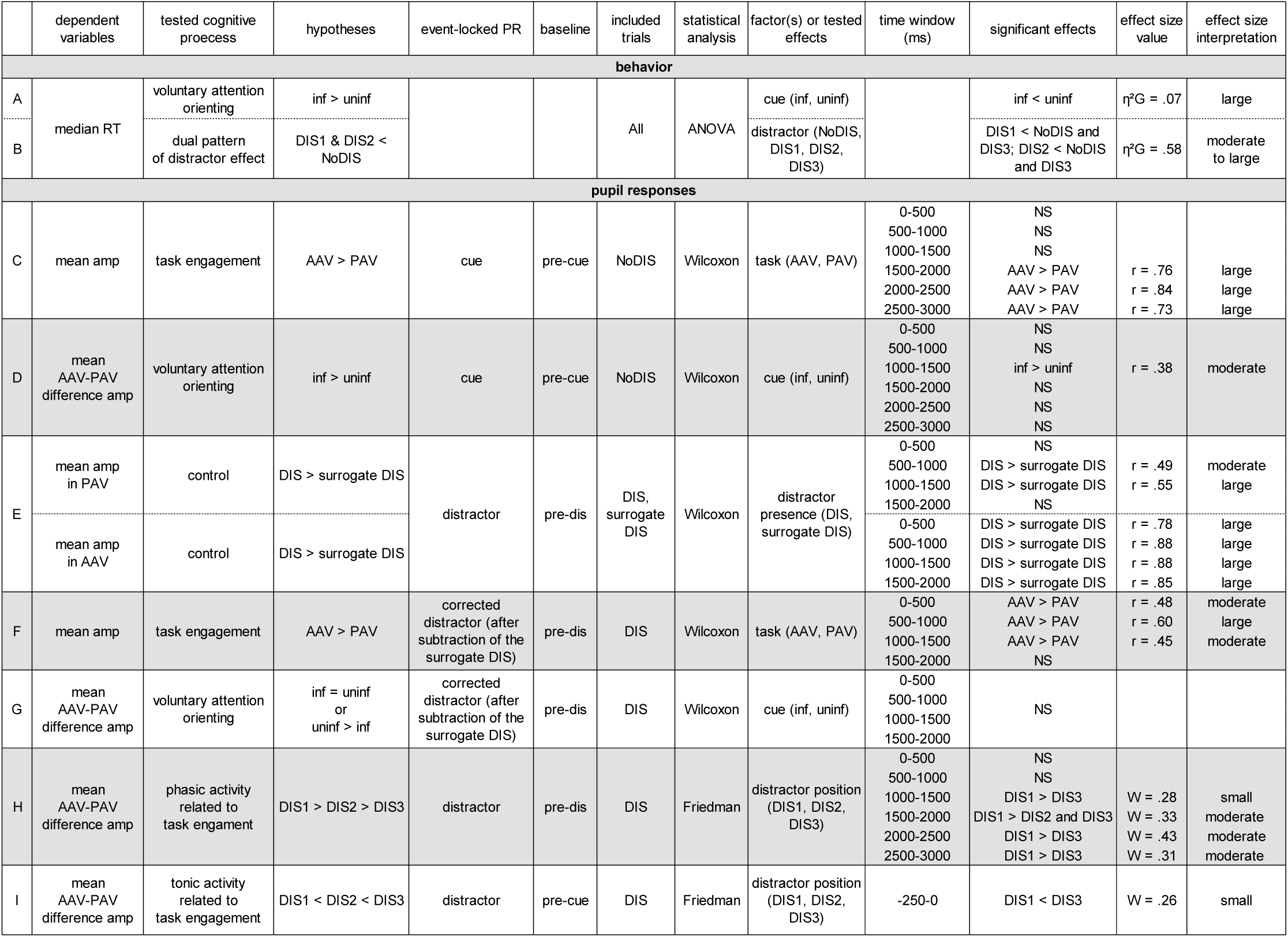
Summary of the statistical analyses and results. For each dependent variable, details and results (significant effect, effect size value and interpretation) of the statistical analyses are provided. The corresponding tested cognitive processes and hypotheses are also provided. RT: reaction time, AAV: Active Audio-Visual condition, PAV: Passive Audio-Visual condition amp: amplitude; inf: informative, uninf: uninformative, DIS: distractor, NS: non-significant.

#### Pupil data

For statistical analysis of the pupil dilation, since data were not normally distributed (all p-value inferior to .05) non-parametric tests were used to determine whether pupil size varies significantly depending on conditions. Wilcoxon signed-rank tests were conducted when only two paired conditions were compared, and Friedman tests were conducted when three paired conditions were compared.

Statistic values (Z for Wilcoxon and X^2^ for Friedman), probability levels and effect sizes (r for Wilcoxon and Kendall’s W for Friedman) for each test are provided. For the Wilcoxon effect size, r value was calculated as Z statistic (extracted from permutation test) divided by the square root of the sample size. For the Friedman test effect size, the Kendall’s W were calculated by dividing the Friedman statistic value X^2^ by the product of the sample size N and one less than the number of measurements per subjects (K-1). Interpretation of r and Kendall’s W uses the Cohen’s interpretation guidelines (Cohen, 1988) of 0.1 to 0.3 (small effect), 0.3 to 0.5 (moderate effect) and >= 0.5 (large effect). Significant effects from Friedman tests were further examined by pairwise comparisons using paired bilateral Wilcoxon signed-rank tests, p-values are adjusted using the Bonferroni multiple testing correction method.

When non-significant effects were found in all tested time windows, Bayesian non-parametric Wilcoxon signed-rank test were conducted using JASP software (JASP—A Fresh Way to Do Statistics, 2021; Version 0.14.1). In contrast to Frequentist statistics, Bayesian analyses allow one to assess the credibility of both the alternative and null hypotheses. We reported Bayes factor (BF10) as a measure of evidence in favor of the null hypothesis (BF_10_ 0.1–0.33, 0.01–0.1, and lower than 0.01: moderate, strong, and decisive evidence, respectively) and in favor of the alternative hypothesis (value of 3–10, 10–100, and more than 100: moderate, strong, and decisive evidence, respectively, Lee and Wagenmakers, 2014).

##### PR to cues, without distracting sounds

Statistical analysis of the cue-locked PR (NoDIS trials) was conducted on the mean PR amplitude in the 0-500ms, 500-1000ms, 1000-1500ms, 1500-2000ms, 2000-2500ms and 2500-3000ms time windows following the cue.

First, to test the task effect, PR amplitudes in the AAV and PAV conditions were compared using paired unilateral Wilcoxon signed-rank tests (AAV > PAV) (Table 1.C).

Second, to test the cue effect on the task engagement, mean difference in PR amplitude between AAV and PAV conditions (AAV – PAV) were computed in the informative and uninformative conditions and compared using paired unilateral Wilcoxon signed-rank tests (Informative > Uninformative) (Table 1.D).

##### PR to distractors: task and cue effects

Statistical analyses of the DIS-locked PR were conducted on the mean PR amplitude in the 0-500ms, 500-1000ms, 1000-1500ms and 1500-2000ms time windows following the distractor.

First, to test the existence of a DIS-locked PR, in the PAV and AAV conditions, the mean amplitude of the DIS-locked PR was compared to the surrogate DIS-locked PR using an unilateral Wilcoxon signed-rank test (DIS-locked > surrogate DIS-locked) (Table 1.E).

Second, to test the task effect, DIS-locked-corrected PR (after subtraction of the surrogate DIS-locked PR) amplitudes in the AAV and PAV conditions were compared using paired unilateral Wilcoxon signed-rank tests (AAV > PAV) (Table 1.F).

Third, to test the cue effect on the task engagement, mean difference in DIS-locked-corrected PR amplitude between AAV and PAV conditions (AAV – PAV) were computed in the informative and uninformative conditions and compared using paired bilateral Wilcoxon signed-rank tests (Table 1.G).

This 3-step analysis was also performed time-locked to cue-onset, comparing DIS and NoDIS trials, in order to ensure that the observed effects are not attributable to differences in baseline amplitude prior to distractor onset (see Supplementary Data for more details).

##### PR to distractors: position effects

To test the distractor position effect, mean difference in DIS-locked PR amplitude (pre-dis baseline correction) between AAV and PAV conditions (AAV – PAV) in the DIS1, DIS2, and DIS3 conditions were computed in consecutive 500 ms wide time windows (i.e., 0-500ms, 500-1000ms, 1000-1500ms, 1500-2000ms, 2000-2500ms and 2500-3000ms) following a distractor and compared using a Friedman test (Table 1.H).

Because pupil size immediately preceding the distractor might depend on the timing of the distractor’s appearance relative to the cue and target, statistical analyses were conducted on the mean pupil size amplitude over the -250 to 0ms period preceding distractor after pre-cue baseline correction (-250 to 0 ms). The mean differences in amplitude between AAV and PAV conditions (AAV – PAV) in the DIS1, DIS2, and DIS3 conditions were compared using a Friedman test (Table 1.I).

## Results

### Behavior

Participants correctly performed the AAV blocks with (mean ± standard deviation) 93.7% ± 2.62% of hits, 3.4% ± 1.4% of late responses, and 2.8% ± 2% of premature responses.

### Reaction times

RTs were significantly shorter for informative than for uninformative cues, as shown by a significant main cue effect (F_(1, 19)_ = 8.48, *p* =.009, η²G = .07) (Figure 2 and Table 1.A).

**Figure 2.**
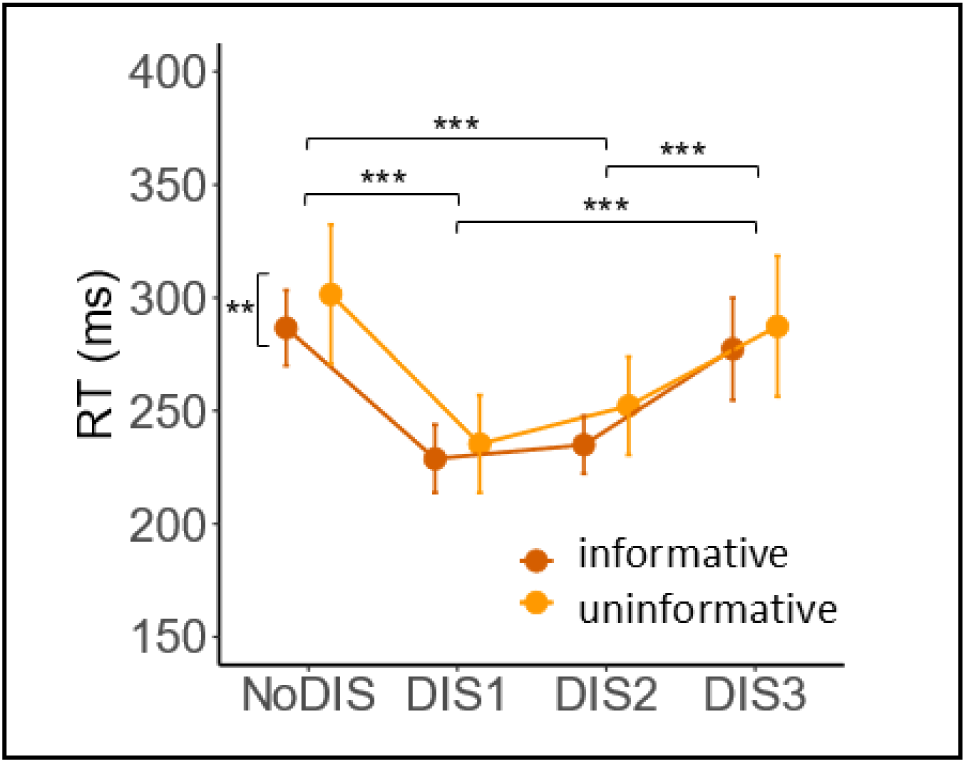
Behavioral results. Mean reaction time as a function of cue type (informative in brown or uninformative in orange) and distractor condition (NoDIS, DIS1, DIS2, DIS3). Error bars represent within-subject standard errors of the mean. ** *p* < .01, *** *p* < .001 (after Bonferroni correction).

A significant main effect of distractor (F_(3, 57)_ = 61.64, *p* = 6.88 x 10^-18^, η²G = .58) was also observed. Pairwise comparisons showed that the RTs were significantly shorter in DIS1 than NoDIS and DIS3 (all *p* < .0001) and in DIS2 than NoDIS and DIS3 (all *p* < .0001). No significant cue by distractor interaction was found (*p* = .729). (Figure 2 and Table 1.B).

### Pupil Responses

In the PAV condition and in the absence of distracting sound, from the cue-locked PR, we first observe a large pupil constriction, also visible in the PV but not the PA condition (see SupFigure1). This constriction can be attributed to the pupillary light response to the cue, which is brighter than the fixation cross (for review see Mathôt, 2018). Then we observe a small pupil dilation, corresponding to the response to the “target” sound, also visible in the PA but not the PV condition.

### PR to cues, without distracting sounds

The pupil dilation response after the relevant cue is larger in amplitude in the active condition compared to the passive condition. This task effect on the cue-locked PR amplitude is larger after an informative rather than an uninformative cue.

To test the task effect on the cue-locked PR, mean amplitudes of the PR in trials with no distracting sound were compared in the PAV and AAV conditions. From 1500 to 3000ms post cue (i.e., in three consecutive 500-ms time windows), the mean amplitude of the cue-locked PR was found larger in the active than in the passive condition (1500-2000 ms: Z = 196, *p* = .0001, r = .76; 2000-2500ms: Z = 206, *p* = 6.7 x 10^-6^, r = .84; 2500-3000ms: Z = 193, *p* = .0002, r = .73) (Figure 3A and Table 1.C). No significant effect of the task was found on the other time-windows (All *p* > .10).

**Figure 3.**
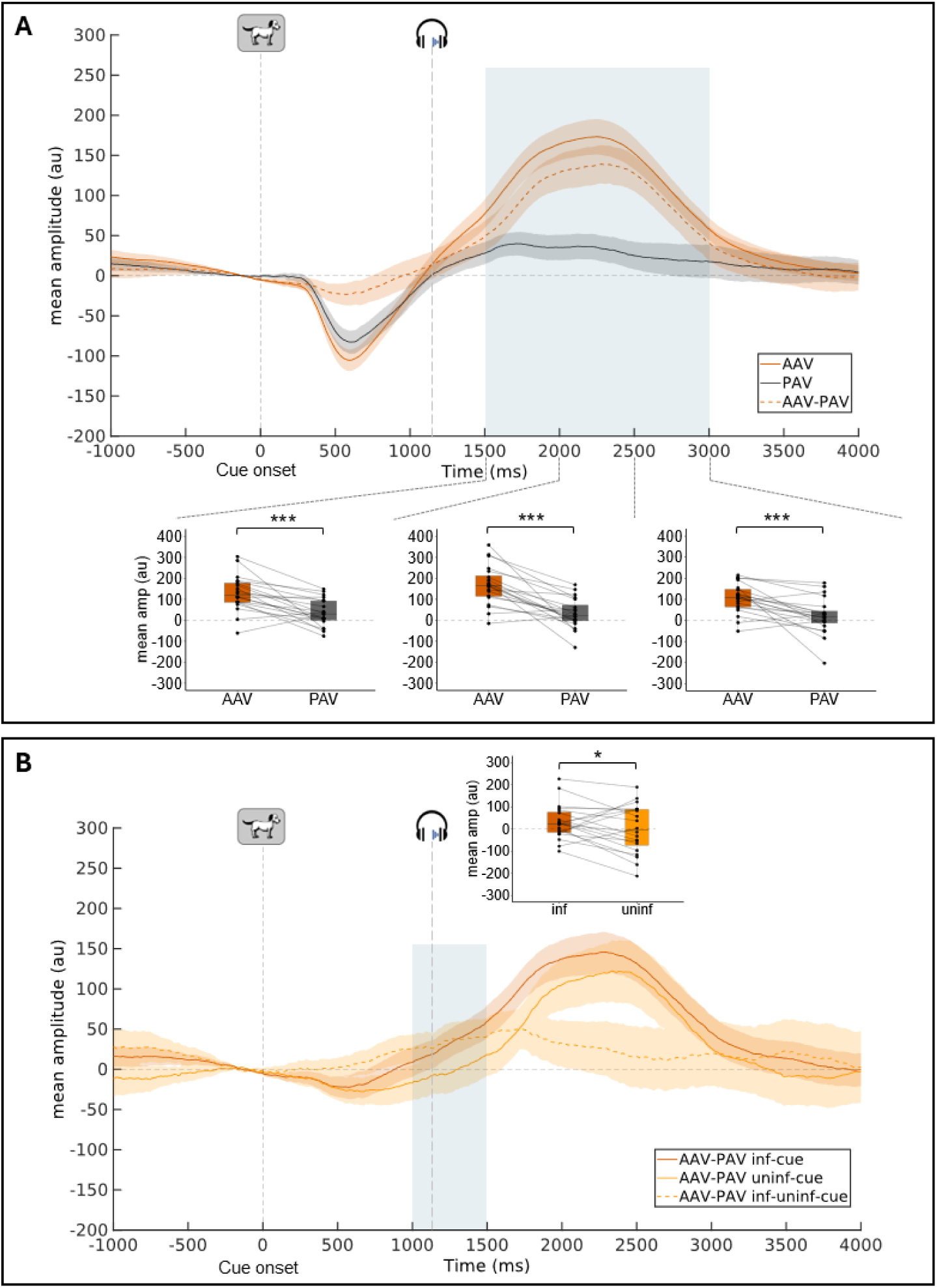
Cue-locked pupil response (group-average, 250-ms pre-cue baseline subtraction) in trials without distractor (NoDIS trials). **A:** Mean pupil dilation in AAV, PAV conditions and the subtraction between the two. **B:** Mean pupil difference curves (AAV-PAV) for informative & uninformative cue conditions. Shadowed areas surrounding the curves represent standard errors of the mean. Examples of stimuli presented to participants are shown at their relative onset (for the target, the mean onset latency is indicated). Blue areas indicate time-windows where the task (A) or the cue (B) effect are significant. For these time windows, boxplots with individual data are depicted (mean pupil dilation amplitude in 500ms time-windows). Within each boxplot, the horizontal line represents the group median, the box the first and third quartiles, the whiskers the largest value under 1.5*IQR. (IQR = inter-quartile range). Superimposed to each boxplot, the dots represent individual means. * *p* < .05, *** *p* < .001. AAV: Active Audio-Visual condition, PAV: Passive Audio-Visual condition, inf: informative, uninf: uninformative.

Then, to test the cue effect on the cue-locked PR, the mean differences in amplitude between active and passive conditions (AAV-PAV) in trials with no distracting sound, were compared in the informative and uninformative cue conditions. From 1000 to 1500ms, the mean AAV-PAV difference in amplitude was larger after an informative than after an uninformative cue (Z = 60, *p* = .048, r = .38) (Figure 3B and Table 1.D). No significant effect of the cue was found on the other time-windows (0-500ms: Z = 107, *p* = .536; 500-1000ms: Z = 78, *p* = .165; 1500-2000ms: Z = 64, *p* = .066; 2000-2500ms: Z = 80, *p* = .184; 2500-3000ms: Z = 88, *p* = .273).

### PR to distractors

#### PR to distractors: task and cue effects

The pupil dilation response to the task-irrelevant distractors is larger in amplitude in the active condition compared to the passive condition. This task effect on the DIS-locked-corrected PR amplitude is not modulated by the cue conditions (informative vs. uninformative).

First, to assess the presence of a PR to distractors in the PAV and AAV conditions, the mean amplitude of the distractor-locked PR was compared to the surrogate DIS-locked PR (same timing between cue and target in trials with no distracting sound). In the PAV condition (see SupFigure2A and Table 1.E), from 500 to 1500ms, the mean amplitude was larger in the presence of distractor (500-1000ms: Z = 164, *p < .05*, r = .49; 1000-1500ms: Z = 171, *p < .01*, r = .55). No significant effect was found on the other time-windows (0-500ms: Z = 87, *p* = .751; 1500-2000ms: Z = 148, *p* = .057). In the AAV condition (see SupFigure2B and Table 1.E), from 0 to 2000ms, the mean amplitude was larger in the presence of distractor (0-500ms: Z = 198, *p* < .001, r = .78; 500-1000ms: Z = 210, *p* < .0001, r = .88; 1000-1500ms: Z = 210, *p < .0001*, r = .88; 1500-2000ms: Z = 207, *p* < .0001, r = .85). A pupil dilation response to distractors is present in both the passive and active conditions.

Second, to test the task effect on the DIS-locked-corrected PR, the mean amplitudes of the PR were compared in the PAV and AAV conditions. From 0 to 1500ms, the mean amplitude of the DIS-locked-corrected PR was larger in the active than the passive condition (0-500ms: Z = 163, *p* = .015, r = .48; 500-1000ms: Z = 177, *p* = .003, r = .60; 1000-1500ms: Z = 159, *p* < .022, r = .45) (Figure 4A and Table 1.F). No significant effect of the task was found on the other time-windows (1500-2000ms: Z = 149, *p* = .053).

**Figure 4.**
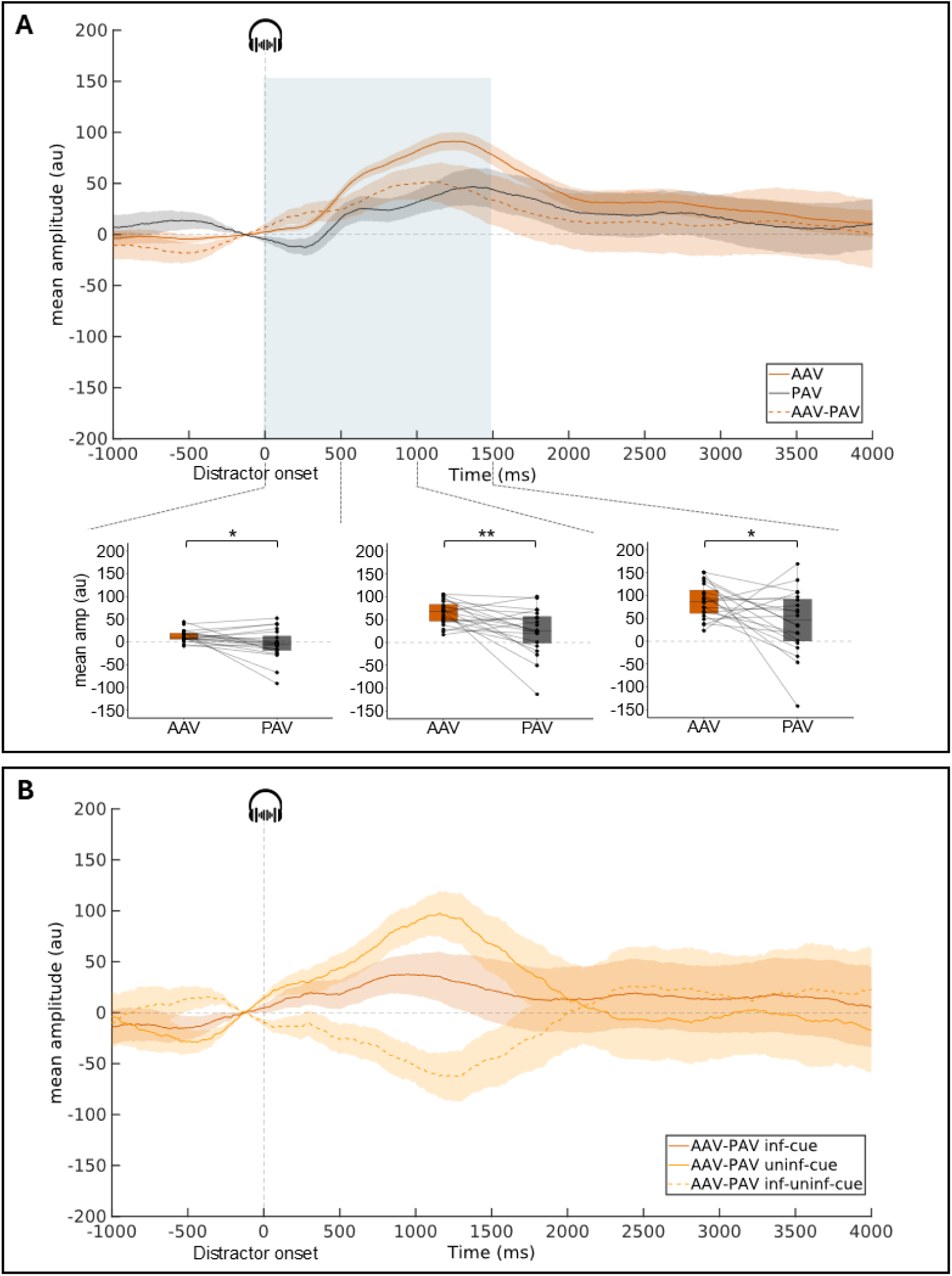
Distractor-locked-corrected pupil response (group-average, subtraction of the surrogate DIS-locked PR & 250-ms pre-distractor baseline subtraction). **A:** Mean pupil dilation in AAV, PAV conditions and the subtraction between the two. **B:** Mean pupil difference curves for informative & uninformative cue conditions. Shadowed areas surrounding the curves represent standard errors of the mean. The blue area (A) corresponds to the time windows where the task effect is significant. For these time windows, boxplots with individual data are depicted (mean pupil dilation amplitude in 500ms time-windows). Within each boxplot, the horizontal line represents the group median, the box the first and third quartiles, the whiskers the largest value under 1.5*IQR. (IQR = inter-quartile range). Superimposed to each boxplot, the dots represent individual means. * *p* < .05, ** *p* < .01. AAV: Active Audio-Visual condition, PAV: Passive Audio-Visual condition, inf: informative, uninf: uninformative.

Third, to test the cue effect on the DIS-locked-corrected PR, the mean differences in amplitude between active and passive conditions (AAV-PAV) were compared in the informative and uninformative cue conditions. No significant cue effect was found on this task difference (*p* > .13 for the 4 time-windows) (Figure 4B and Table 1.G). Bayesian Wilcoxon tests showed no conclusive evidence for or against a cue effect from 0 to 1500 ms (0.371 < BF_10_ < 1.185) and positive evidence for no cue effect on the DIS-locked-corrected PR from 1500 to 2000 ms (BF_10_ = 0.277).

This 3-step analysis was also performed time-locked to cue-onset, comparing DIS and NoDIS trials, and yielded highly similar results (see Supplementary Data and Figures 3 & 4 for more details).

#### PR to distractors: position effects

The task effect on the DIS-locked PR amplitude is larger after an early than a late distractor.

To test the distractor position effect on the DIS-locked PR, the mean differences in amplitude between active and passive conditions (AAV-PAV) were compared. A significant effect of the distractor position was found between 1000 and 3000ms post-distractor onset, i.e., in four consecutive 500-ms time windows (1000-1500ms: X^2^ = 11.1, *p* = .004, W = .28; 1500-2000ms: X^2^ = 13.3, *p* = .001, W = .33; 2000-2500ms: X^2^ = 17.2, *p* < .001, W = .43; 2500-3000ms: X^2^ = 12.4, *p* < .01, W = .31; Figure 5A and Table 1.H). No significant effect of the distractor position was found in the other time-windows (*p* > .12). From 1000 to 3000ms, pairwise Wilcoxon signed rank comparisons showed that the difference of mean amplitudes between active and passive conditions was significantly larger after DIS1 than DIS2 from 1500-2000ms (Z = 172, *p* < .05) and after DIS1 than DIS3 from 1000 to 3000ms (1000-1500ms: Z = 186, *p* < .01; 1500-2000ms: Z = 204, *p* < .0001; 2000-2500ms: Z = 210, *p* < .0001; 2500-3000ms: Z = 193, *p* < .001).

**Figure 5.**
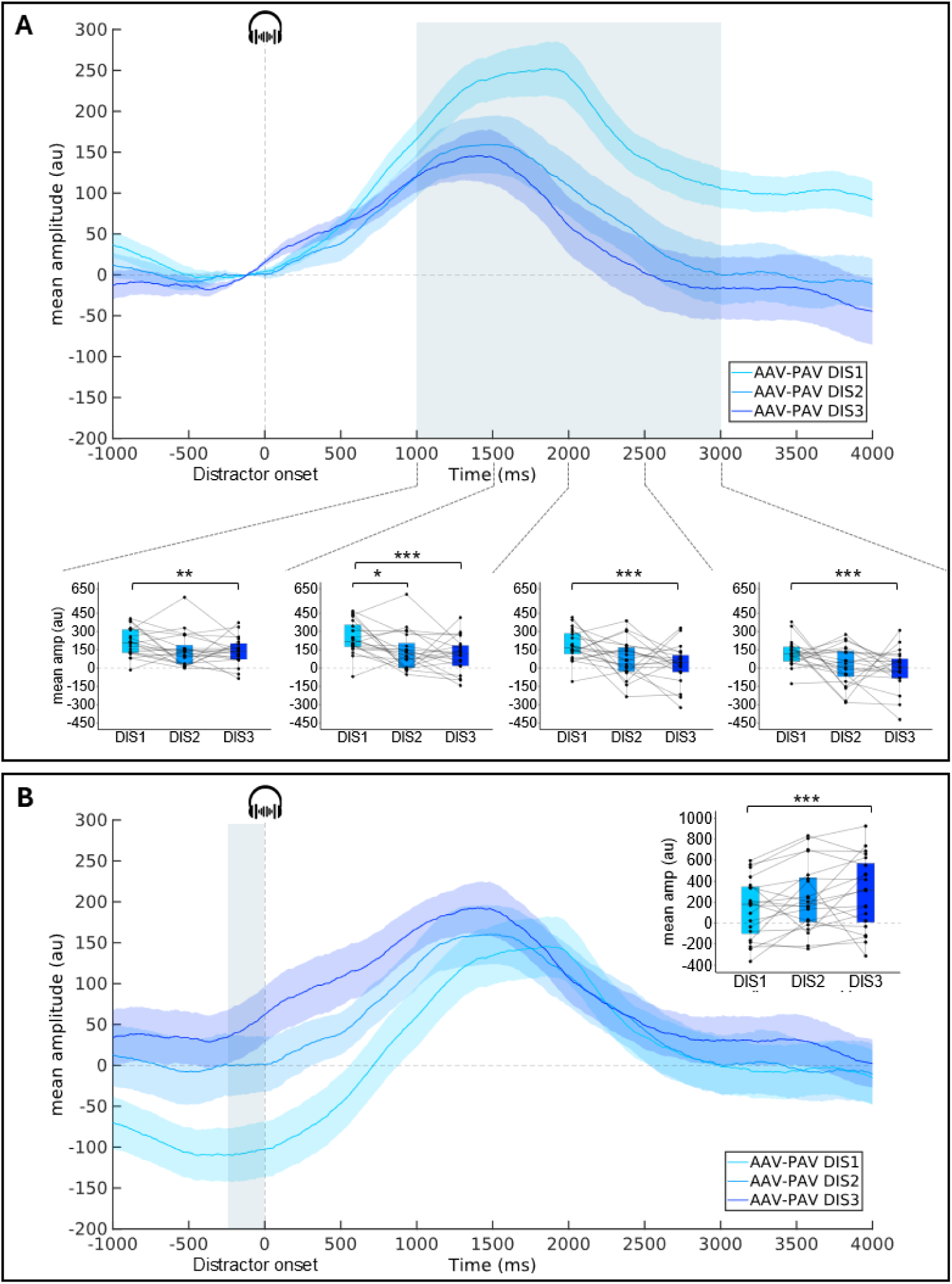
Distractor-locked pupil response (group-average, AAV-PAV subtraction curves) as a function of the distractor position (DIS1, DIS2, DIS3). **A**: with a 250-ms pre-distractor baseline subtraction. **B**: with a 250-ms pre-cue baseline correction. Shadowed areas surrounding the curves represent standard errors of the mean. Blue areas correspond to the time window where the distractor position is significant. For these time windows, boxplots with individual data are depicted (mean pupil dilation amplitude in 500ms time-windows). Within each boxplot, the horizontal line represents the group median, the box the first and third quartiles, the whiskers the largest value under 1.5*IQR. (IQR = inter-quartile range). Superimposed to each boxplot, the dots represent individual means. * *p* < .05, ** *p* < .01, *** *p* < .001. AAV-PAV: Active Audio-Visual - Passive Audio-Visual condition.

The task effect on the pre-distractor pupil size is lower before an early than a late distractor.

Finally, to test the distractor position effect on the pre-distractor pupil size, the mean differences in amplitude between active and passive conditions (AAV-PAV) were compared during the 250ms preceding the distractor onset (-250-0ms) after pre-cue baseline correction. A significant effect of the distractor position was found (X^2^ = 10.3, *p* = .01, W = .26; Figure 5B and Table 1.I). Pairwise Wilcoxon signed rank test showed that the difference of baseline between active and passive conditions was lower during the 250ms preceding DIS1 than DIS3 (Z= 16, *p* < .001). No other significant effect was found (All *p* > .13).

## Discussion

Here, we used pupillometric responses recorded during the Competitive Attention Test (CAT, Bidet-Caulet et al., 2015; Masson et Bidet-Caulet, 2019; Hoyer et al., 2021, 2023a, 2023b) to investigate how attention interacts with arousal as a function of task engagement and stimulus relevance. To explore task engagement, we compared pupil dilation in participants who either actively performed the CAT or were passively presented with the same sequences of stimuli. In addition, because the CAT incorporates both informative and uninformative cues, it allows to examine how voluntary attention orienting further modulates the pupil responses (PR) to relevant versus irrelevant sounds. By employing pupillometry in the CAT, we thus aimed to directly assess whether NE-driven arousal and attention interact both globally and selectively way.

Behavioral results in the active condition closely mirrored earlier findings using a slightly different version of the CAT (e.g., Bidet-Caulet et al., 2015; Hoyer et al., 2023a, 2023b). Specifically, participants exhibited shorter reaction times when cues were informative rather than uninformative (i.e., cue effect), indicating stronger voluntary attention orienting (Table 1.A). Moreover, although the presence of an early distracting sound speeded up responses, reaction times (RT) lengthened as the distractor appeared closer to the target (Table 1.B). This dual pattern replicates previous CAT results, reflecting both a phasic arousal benefit (shorter RT) and a distraction cost (longer RT, Bidet-Caulet et al., 2015; Masson & Bidet-Caulet, 2019; Hoyer at al., 2021, 2023a, 2023b). Hence, this continued responsiveness to irrelevant distractors, under active engagement, suggests a nuanced interplay between globally elevated arousal and attention to distractors, prompting a deeper pupillometric investigation of these processes.

In the passive condition, we observed an initial pupil constriction in response to the bright visual cue, followed by a dilation in anticipation of - and in response to - the target sound, as well as in response to distracting sounds. These passive-condition findings indicate that both target and distracting sounds in the CAT elicit phasic arousal, even in the absence of an active task. By contrast, under active task condition, both relevant (target) and irrelevant (distractor) sounds elicited a significantly larger increase in pupil dilation response (Table 1.C, F). This finding aligns with previous work showing that greater task engagement - through heightened difficulty, multitasking, or active vs. passive comparisons - yields larger pupil dilation responses (Kahneman & Beatty, 1967; Kramer et al., 2012; Lisi et al., 2015; Laeng et al., 2016; Liao et al., 2016). Indeed, actively engaging in a task involves focusing attentional resources (Tops and Boksem, 2010), which increases phasic arousal (Kahneman, 1973; Posner and Peterson, 1990; Aston-Jones and Cohen, 2005), as reflected by increased pupil dilation. Such elevated arousal in response to both relevant and irrelevant events suggests a global, transient boost in brain excitability arising from task engagement, largely independent of stimulus relevance. Hence, these findings indicate that the global rise in arousal from active task engagement extends to both relevant and irrelevant stimuli.

Moreover, our results revealed a larger pupil dilation 1000 to 1500 ms post-cue in the informative compared to the uninformative condition (Table 1.D). Because this rise begins just before the target, it likely reflects both anticipatory and response-related processes for the relevant target sound. Another study using the CAT paradigm showed the same pattern suggesting that, despite the modest sample size, this effect is robust and replicable (Grandjean et al., 2024). This finding aligns with the limited literature examining how voluntary attention orienting affects pupil dilation (Dragone et al., 2018; but see Aminihajibashi et al., 2020). Notably, this pupil dilation starts before target onset, indicating a transient phasic arousal boost driven by top-down attention mechanisms during target expectation (Totah et al., 2021; Aston-Jones & Cohen, 2005). In contrast, the pupil dilation response to the irrelevant distractor was unaffected by whether the cue was informative or uninformative (Table 1.G). Bayesian analyses further indicated a similar pupil response to distractor after both cue types, suggesting that voluntary orienting does not modulate the phasic response to irrelevant events. Thus, stronger voluntary attention orienting selectively increases phasic arousal to relevant stimuli. These findings thus underly the selective nature of arousal modulations, demonstrating that voluntary orienting specifically enhances phasic responses to relevant stimuli while leaving irrelevant events unaffected.

Interestingly, we also found a larger PR to task-irrelevant distractors when they were presented long before the target, compared to those presented just before the target sound (Table 1.H). Two possible explanations might account for this result. First, the larger PR to early distracting sounds may result from more overlap with pupil constriction in response to the cue and from less temporal overlap with the subsequent target-locked response. Second, as attentional and motor preparation ramp up closer to the target onset, the pupil response to irrelevant sounds may diminish as attentional and motor preparation increase. Indeed, previous studies have shown that longer cue-to-target delays improve preparation (Posner et al., 1980; Niemi & Näätänen, 1981; Coull & Nobre, 1998; Hackley & Valle-Inclán, 1999; Griffin et al., 2001) and performance in detection tasks (Griffin et al., 2001). Thus, as the target approaches, top-down mechanisms – such as enhanced voluntary attentional orienting and inhibition – intensify, likely reducing the phasic arousal response to irrelevant events. This latter account aligns with our observation of increased pre-distractor baseline pupil dilation with approaching target onset (Table 1.I). Such baseline pupil changes have been related to tonic activity in the locus coeruleus (LC), which reflects task engagement and sustained tonic arousal (Aston-Jones & Cohen, 2005; Gilzenrat et al., 2010; Jepma & Nieuwenhuis, 2011; Murphy et al., 2014; Unsworth & Robison, 2016, 2018; Konishi et al., 2017). Overall, these findings show that increased top-down attention during target anticipation can be captured by shifts in baseline pupil size (i.e. across the cue-target delay). Collectively, the timing-dependent variation in distractor-evoked pupil responses indicates that phasic arousal to irrelevant sounds wanes as top-down target preparation intensifies, suggesting that the global boost from task engagement is modulated by task timing rather than applied uniformly.

We observed that heightened arousal response to auditory stimuli is mediated by task engagement (active vs. passive), independently from the stimulus relevance. We found that manipulating voluntary attention orienting with informative and uninformative cues selectively increased arousal for relevant target sounds. Taken together, our findings align with the Glutamate Amplifies Noradrenergic Effects (GANE) model (Mather et al. 2016), which posits that an NE-mediated rise in arousal amplifies high-priority neural representations while suppressing lower-priority representations, thereby enhancing selectivity. Therefore, top-down attention might selectively engage LC-NE phasic activity based on stimulus relevance, enabling the LC-NE system to bolster selective attention (Dahl et al., 2022) by enhancing cortical excitability for relevant stimuli. In other words, the present data suggest the implication of the LC-NE system in selective attention.

## Conclusion

By simultaneously manipulating task engagement (active vs. passive) and stimulus relevance (informative vs. uninformative cues) in a cued auditory detection task, the Competitive Attention Test, we examine how attention interacts with arousal. Our results confirm that active task engagement is associated with a global enhancement in pupil-based arousal, while voluntary attention is linked to a selective increase in arousal for high-priority targets. Thus, norepinephrine’s dual role in boosting overall alertness and amplifying relevant information, as proposed by the GANE model, is evident within a single paradigm, advancing our understanding of how attention and arousal intersect and offering insights for optimizing both research designs and applied settings. Beyond its theoretical significance, this study shows that the CAT also incorporates measurable arousal components. This finding further supports the CAT’s relevance for assessing clinical populations, such as those with neurodevelopmental (e.g., ADHD) or neurodegenerative (e.g., Parkinson’s disease) disorders, where attention and arousal mechanisms can both be compromised.

## Supporting information

SupFig

SupData

## Declarations

### Ethics approval statement

The experimental paradigm was approved by a French ethical committee, Comité de Protection des Personnes Sud-Est IV, number 11/90, authorization B11291-10. All participants gave written informed consent (according to the Declaration of Helsinki).

### Data accessibility statement

The dataset supporting the conclusions of this article will be available on OSF: https://osf.io/8msfw/overview.

### Competing interests

The authors declare that they have no competing interests.

### Authors’ contributions

R.S.H., A.M. and A.B-C. designed research; R.S.H. acquired data; A.M., A.B-C., R.S.H. and A.G. wrote the codes for data analysis; A.G. analyzed data; A.G. and A.B-C. wrote the paper; R.S.H., A.C., A.M. and A.B-C. edited the paper.

## Funding

This work was supported by two grants from the French National Research Agency (ANR) awarded to A. Bidet-Caulet (ANR-14-CE30-0001-01 & ANR-19-FRAL-0007-01). This work was performed within the framework of the LABEX CORTEX (ANR-11-LABX-0042) and the LABEX CeLyA (ANR-10-LABX-0060) of Université de Lyon, within the program “Investissements d’Avenir” (ANR-16-IDEX-0005) operated by the French ANR.

## Acknowledgements

The authors thank M. Gaudet-Trafit for his help with data preprocessing and analysis. We also would like to thank all participants.

## Notes

### Competing Interest Statement

The authors have declared no competing interest.

### Summary of Updates

We re-analyzed the data time-locked to cue onset and compared trials with and without distractors. The results were highly consistent with those obtained when time-locked to distractor onset. These additional analyses have been incorporated into the manuscript as Supplementary Data.

