## Supplementary material for "Global and selective effects of auditory attention on arousal: insights from pupil dilation": SupFig

### Supplementary figures

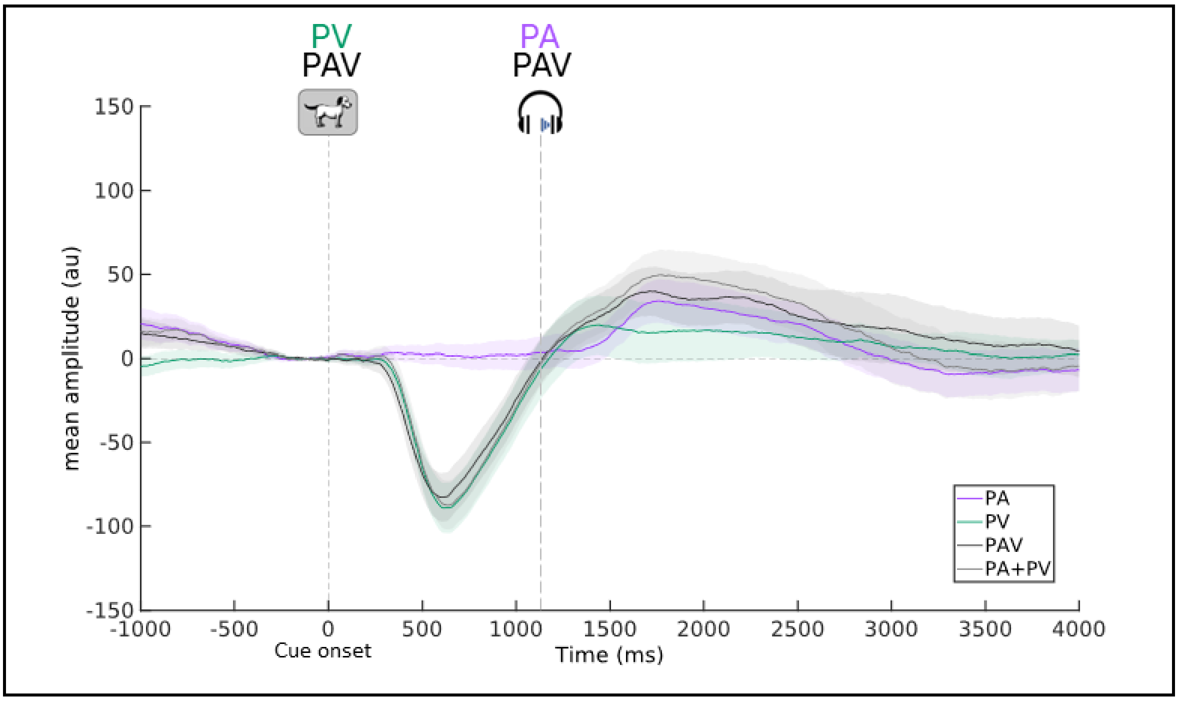


**Sup Figure 1. Cue-related pupil response** (group-average, 250-ms pre-cue baseline subtraction) in NoDIS trials, in Passive Audio (PA, purple line), Passive Visual (PV, green line), the sum of Passive Audio + Passive Visual (PA+PV, grey line) and Passive Audio-Visual (PAV, black line) conditions. Stimuli presented to participants according to conditions are shown at their relative onset (for the target, the mean onset latency is indicated). It is interesting to note the PDR additivity with the PDR in PAV corresponding to the sum of PDRs in PA and PV.

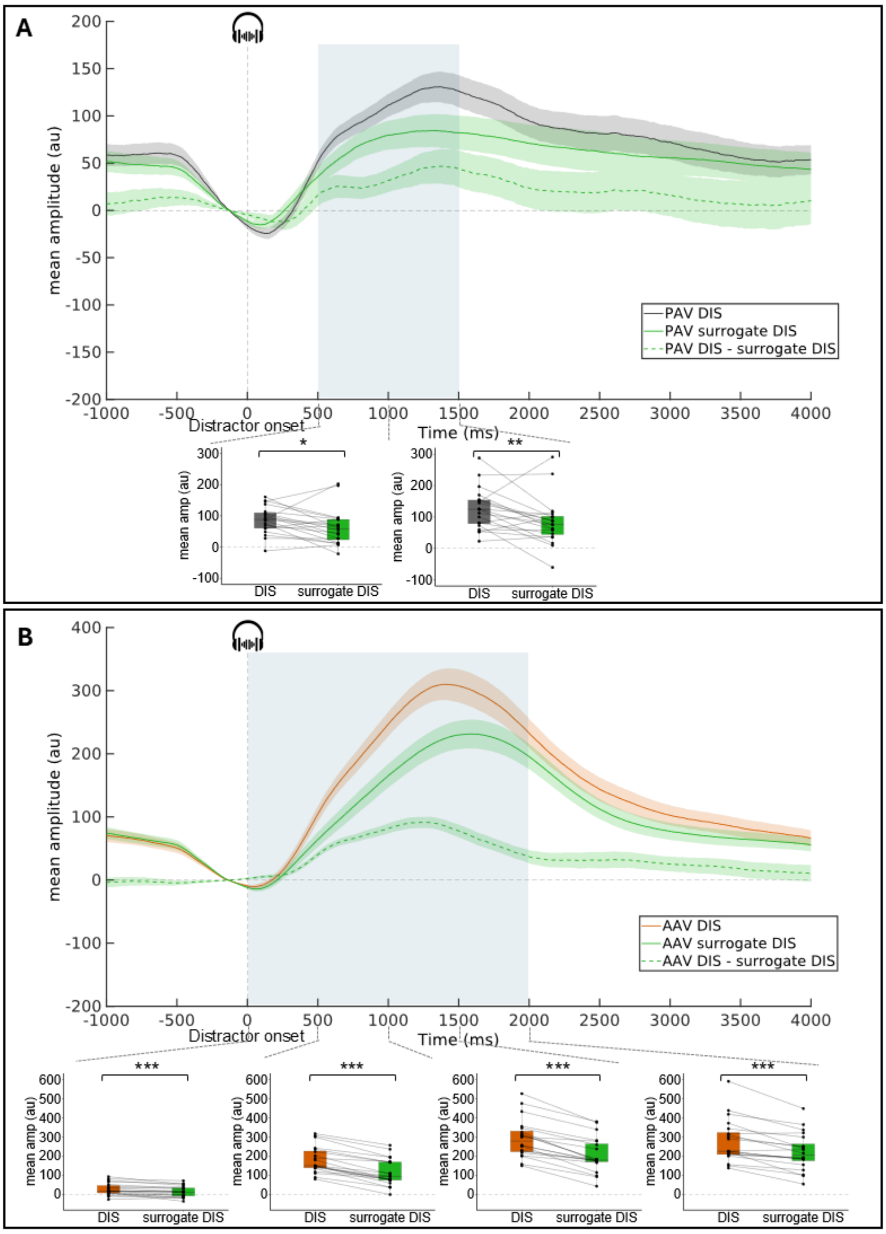


**Sup Figure 2. Distractor-locked pupil response** (group-average, subtraction of the surrogate DIS-locked PR & 250-ms pre-distractor baseline subtraction) **in trials with (DIS trials), without distractor (NoDIS trials) and the subtraction between the two. A:** Mean pupil dilation in PAV. **B:** Mean pupil dilation in AAV. Shadowed areas surrounding the curves represent standard errors of the mean. Blue areas indicate time-windows where the distractor effect is significant. For these time windows, boxplots with individual data are depicted (mean pupil dilation amplitude in 500ms time-windows). Within each boxplot, the horizontal line represents the group median, the box the first and third quartiles, the whiskers the largest value under 1.5*IQR. (IQR = inter-quartile range). Superimposed to each boxplot, the dots represent individual means. * $p < .05$, ** $p < .01$, *** $p < .001$. AAV: Active Audio-Visual condition, PAV: Passive Audio-Visual condition, DIS: distractor.

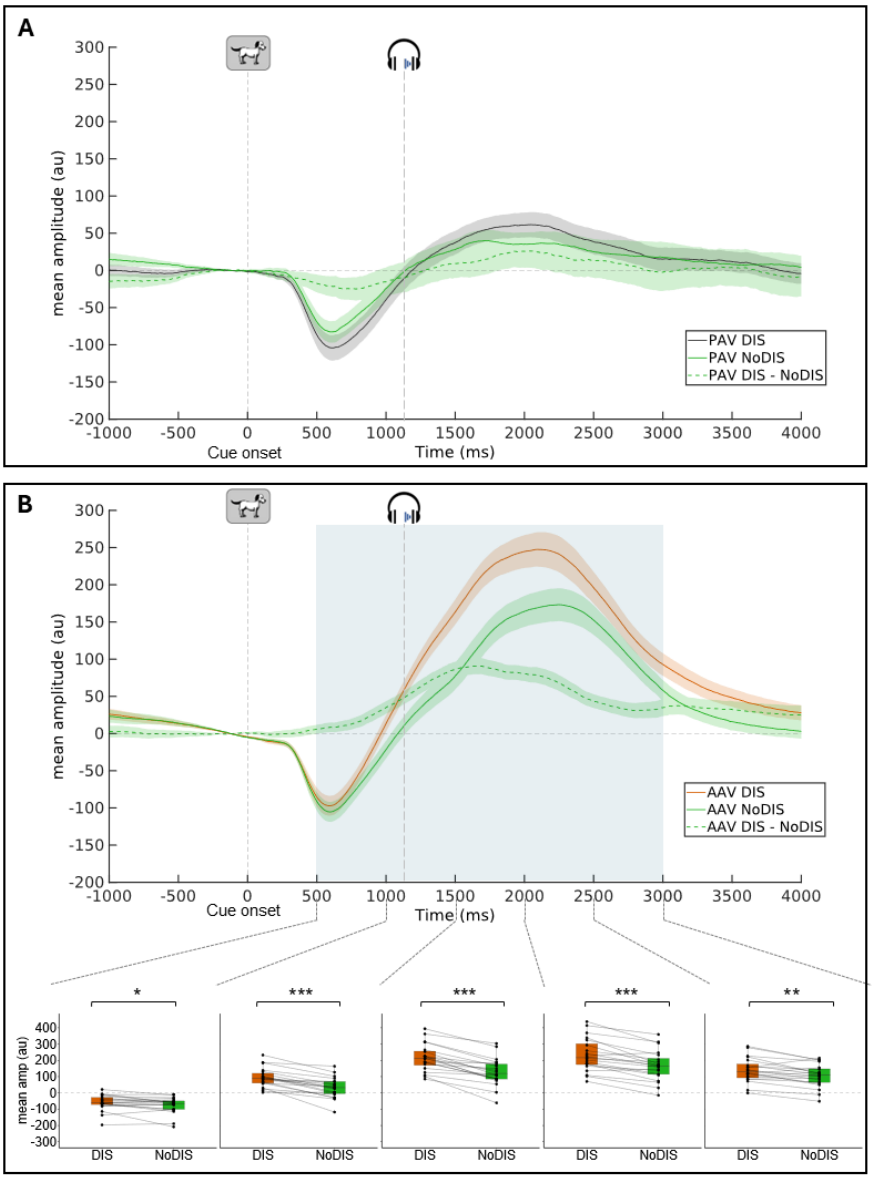


**Sup Figure 3. Cue-locked pupil response** (group-average, 250-ms pre-cue baseline subtraction) **in trials with (DIS trials), without distractor (NoDIS trials) and the subtraction between the two. A:** Mean pupil dilation in PAV. **B:** Mean pupil dilation in AAV. Shadowed areas surrounding the curves represent standard errors of the mean. Examples of stimuli presented to participants are shown at their relative onset (for the target, the mean onset latency is indicated). Blue areas indicate time-windows where the distractor effect is significant. For these time windows, boxplots with individual data are depicted (mean pupil dilation amplitude in 500ms time-windows). Within each boxplot, the horizontal line represents the group median, the box the first and third quartiles, the whiskers the largest value under 1.5*IQR. (IQR = inter-quartile range). Superimposed to each boxplot, the dots represent individual means. * p < .05, ** p < .01, *** p < .001. AAV: Active Audio-Visual condition, PAV: Passive Audio-Visual condition, DIS: distractor, NoDIS: no-distractor.

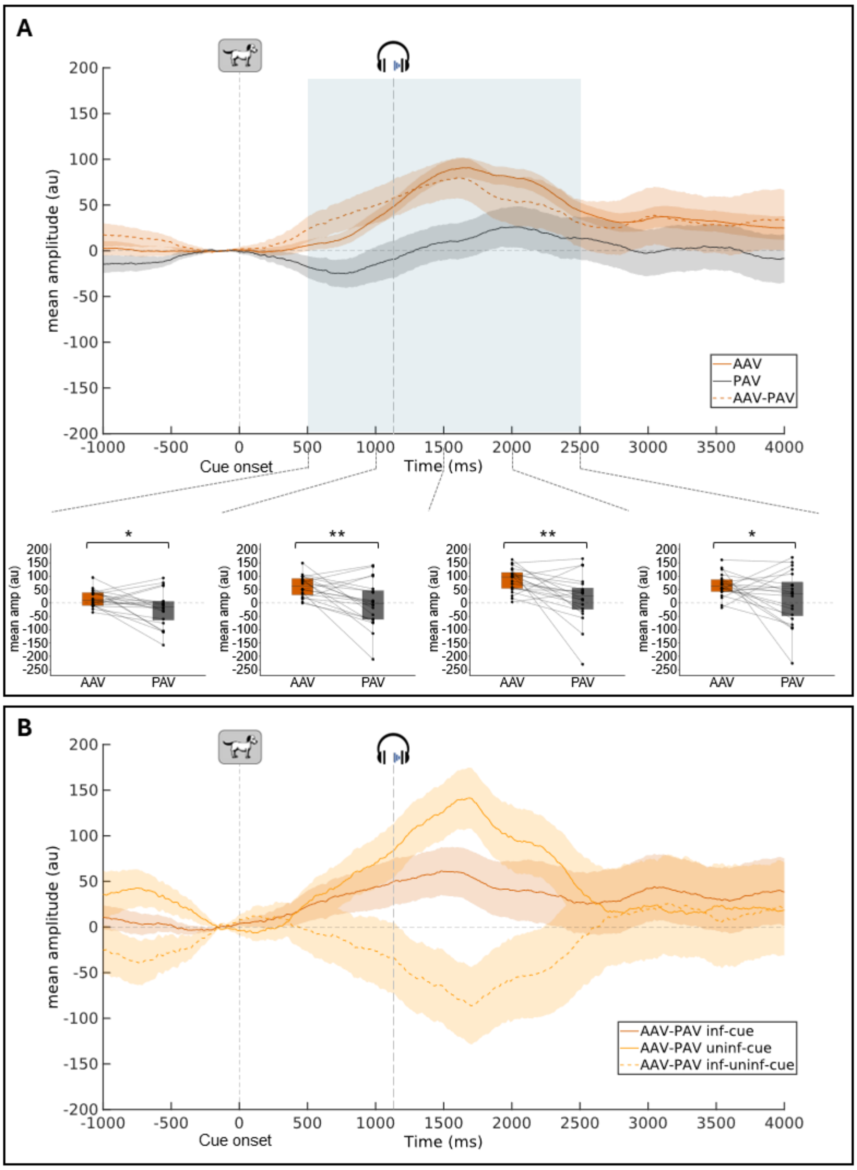


**Sup Figure 4. Cue-locked pupil response** (group-average, 250-ms pre-cue baseline subtraction) **after subtraction between DIS and NoDIS trials. A:** Mean pupil dilation in AAV, PAV conditions and the subtraction between the two. **B:** Mean pupil difference curves (AAV-PAV) for informative & uninformative cue conditions. Shadowed areas surrounding the curves represent standard errors of the mean. Examples of stimuli presented to participants are shown at their relative onset (for the target, the mean onset latency is indicated). Blue areas indicate time-windows where the task (A) or the cue (B) effect are significant. For these time windows, boxplots with individual data are depicted (mean pupil dilation amplitude in 500ms time-windows). Within each boxplot, the horizontal line represents the group median, the box the first and third quartiles, the whiskers the largest value under 1.5*IQR. (IQR = inter-quartile range). Superimposed to each boxplot, the dots represent individual means. * $p < .05$, ** $p < .01$. AAV: Active Audio-Visual condition, PAV: Passive Audio-Visual condition, inf: informative, uninf: uninformative.
