## Supplementary material for "Global and selective effects of auditory attention on arousal: insights from pupil dilation": SupData

### Supplementary data

#### Results section

##### PR to cues, with distracting sounds

To test the distractor effect on the cue-locked PR in the PAV condition, the mean amplitudes of the PDR were compared in the distractor and no-distractor conditions. No significant distractor effect was found in the PAV condition (p > .11 for the six 500-ms time-windows from 0 to 3000 ms) (**Sup Figure 3A**).

To test the distractor effect on the cue-locked PR in the AAV condition, the mean amplitudes of the PR were compared in the distractor and no-distractor conditions. From 500 to 3000ms, the mean amplitude was larger in the distractor than the no-distractor condition (500-1000ms: Z = 153, $p < .05$, r = .40; 1000-1500ms: Z = 209, $p < .0001$, r = .87; 1500-2000ms: Z = 210, $p < .0001$, r = .88; 2000-2500ms: Z = 207, $p < .0001$, r = .85; 2500-3000ms : Z = 179, $p < .01$, r = .62) (**Sup Figure 3B**). No significant effect was found before 500 ms (0-500ms: Z = 104.5, $p = .51$). These data clearly show a pupil dilation response to distracting sounds in the AAV condition.

Then, to test the task effect on the cue-locked PR to distractor, the mean differences in amplitude between distractor and no-distractor conditions (DIS-NoDIS) were compared in the active and passive conditions. From 500 to 2500ms, the mean DIS-NoDIS difference in amplitude was larger in the active than the passive condition (500-1000ms: Z = 154, $p < .05$, r = .41; 1000-1500ms: Z = 175, $p < .01$, r = .58; 1500-2000ms: Z = 182, $p < .01$, r = .64; 2000-2500ms: Z = 154, $p < .05$, r = .41) (**Sup Figure 4A**). No significant effect of the task was found on the other time-windows (0-500ms: Z = 128, $p = .2$; 2500-3000ms: Z = 137, $p = .12$). The pupil dilation response to distractor is larger in the active than in the passive condition.

Finally, to test the cue effect on the cue-locked PR to distractor, the mean differences in amplitude between active and passive conditions (AAV-PAV) of the PR to distractor (DIS-NoDIS) were compared in the informative and uninformative cue conditions. No significant effect of the cue was found on these differences ($p > .14$ for the 6 time-windows) (**Sup Figure 4B**). Bayesian Wilcoxon tests showed no conclusive evidence for or against a cue effect from 1000 to 2000 ms ($0.681 < BF_{10} < 0.746$) and positive evidence for no cue effect from 0 to 1000 ms (0-500ms: $BF_{10} = .248$; 500-1000ms: BF = .323) and 2000 to 3000ms (2000-2500ms: BF = .323; 2500-3000ms: BF = .239). There is no clear evidence for an effect of the cue on the pupil dilation response to task-irrelevant distractors.
